# Lorentz Tangent Diffusion Adapter for Multi-label Prediction of Protein–Protein Interaction Types

**DOI:** 10.64898/2026.09.28.755203

**Authors:** Qingzhi Yu, Xiang Cheng, Zhengrong Xi, Xuan Xiao

## Abstract

Protein–protein interactions shape cellular organization and disease processes, but their prediction remains difficult because known interaction networks are incomplete, noisy and highly uneven across proteins. Hyperbolic graph learning is well suited to the hierarchical structure of such networks, yet most methods emphasize learning hyperbolic embeddings and pay less attention to how these embeddings are propagated and decoded for type prediction. Here we introduce the Lorentz Tangent Diffusion Adapter (LTDA), a post-embedding module for PPI type prediction, that is, for predicting which of several interaction types a candidate protein pair undergoes. LTDA learns relation-specific protein representations on the Lorentz manifold, projects them to tangent space, applies lightweight graph diffusion, and decodes the diffused representations directly in tangent space. This design uses the tangent space as a prediction-oriented interface between hierarchical representation learning and interaction classification. On SHS27K and SHS148K under BFS and DFS splits, LTDA consistently improves over representative PPI methods and a reproduced HI-PPI baseline, with average gains of 1.41 percentage points in Micro-F1, 3.21 in AUPR, 1.31 in AUC and 1.25 in accuracy. Ablation, degree-bin and biological-evidence analyses indicate that the gains are not explained by simple smoothing or hub correction. LTDA therefore shows that post-embedding adaptation can make Lorentz representations more effective for PPI type prediction.

## 1. Introduction

Protein–protein interactions (PPIs) are fundamental to cellular functional regulation and are widely involved in key biological processes such as signal transduction, metabolic regulation, immune response, and protein complex formation [1, 2]. Accurate identification and prediction of PPIs not only help reveal protein functions and cellular regulatory mechanisms, but are also of great importance for disease mechanism analysis, drug target discovery, and functional genomics. However, experimental determination of PPIs is usually costly and time-consuming, and different experimental platforms vary in coverage and reliability [1, 3]. Meanwhile, existing PPI networks still contain a large number of missing edges, noisy edges, and insufficiently annotated proteins. Therefore, how to fully exploit protein sequence features, functional information, and network topology for high-accuracy PPI prediction remains an important problem in computational biology and bioinformatics.

Computational PPI prediction has evolved from feature-engineered methods and traditional machine learning classifiers to deep representation learning and graph-based modeling. Early computational methods mainly relied on manually designed protein descriptors, such as sequence composition, physicochemical properties, evolutionary profiles, domain information, structural similarity, or other similarity measures. These descriptors were commonly combined with traditional machine learning classifiers, including support vector machines, random forests, (k)-nearest neighbors, logistic regression, and ensemble models, to determine whether a candidate protein pair interacts. Although these methods provided useful computational alternatives to experimental screening, their performance was often limited by feature engineering and by the difficulty of capturing complex nonlinear and network-context-dependent relationships between proteins. Deep learning methods have further reduced reliance on handcrafted descriptors by enabling the learning of protein representations from sequences, structures, or pairwise features. With the development of graph representation learning, graph neural networks (GNNs) have been widely used to model topological dependencies in PPI networks [4–9]. By aggregating neighborhood information, GNNs can capture local associations between proteins and have achieved promising performance in interaction prediction tasks. Meanwhile, PPI prediction has advanced beyond binary interaction detection to more informative multi-type and multi-label prediction. Many earlier studies focused on determining whether two proteins interact. However, real PPI networks contain heterogeneous interaction semantics, and the same protein pair may participate in multiple biological processes simultaneously. Therefore, multi-type PPI prediction requires the model not only to identify potential interacting pairs, but also to predict one or more interaction types that may coexist for the same protein pair. In this work we therefore address interaction-type prediction rather than binary interaction detection. This introduces additional challenges, including label imbalance, interaction-type overlap, and stronger dependence on network context. For example, GNN-PPI learns unknown protein interactions from correlation information using graph neural networks, demonstrating the importance of graph structural information for PPI prediction [6]. In addition, comparative studies on different GNN architectures for PPI prediction have also shown that graph structure modeling plays an important role in improving protein interaction prediction performance [9]. Building upon this line of work, methods such as HIGH-PPI, MAPE-PPI, and HI-PPI further incorporate higher-order neighborhoods, structural encoding, and hierarchical information to enhance the representation capability of models for complex PPI networks [10–13]. In particular, HI-PPI introduces hierarchical structure in PPI networks into hyperbolic representation learning and combines it with interaction-specific learning for multi-type PPI prediction [12], highlighting the potential of hyperbolic geometry in modeling the hierarchical organization of PPI networks.

PPI networks are not simple homogeneous graph structures, but exhibit complex topological properties such as hub nodes, functional modules, local communities, and hierarchical organization [14–16]. For networks with hierarchical and scale-free characteristics, local neighborhood aggregation in Euclidean space may be insufficient to fully capture their non-uniform structures [14–16]. Hyperbolic space, due to its exponential volume growth property, is considered more suitable for representing tree-like, hierarchical, and scale-free networks [17–21]. Previous studies have shown that hyperbolic graph neural networks can more effectively embed hierarchical graph structures in non-Euclidean space and achieve favorable performance in node classification, link prediction, and biological network modeling tasks [17, 19, 22–24]. In typical hyperbolic graph learning frameworks, hyperbolic manifolds are used to represent hierarchical structures, while tangent spaces provide locally Euclidean computational interfaces, making linear transformation, neighborhood aggregation, and downstream prediction easier to implement [17–19, 21–24].

Although hyperbolic representation learning has been applied to PPI prediction, existing methods mainly focus on how to embed proteins into hyperbolic space, while paying less systematic attention to the post-embedding stage, namely how the learned hyperbolic representations should be propagated, adapted, and decoded for downstream prediction. Hyperbolic embedding itself does not necessarily guarantee that the hierarchical information encoded in the manifold is fully accessible to the classifier. This issue is particularly important for multi-type PPI prediction, where the model must not only represent proteins in a hierarchical network geometry, but also transform such latent geometric structure into prediction-effective signals for distinguishing different interaction types. Therefore, after hyperbolic graph encoding, the question of how to adapt and decode the resulting representations remains an important but underexplored problem.

This study is motivated by this post-embedding problem in PPI type prediction. In our preliminary ablation analysis, we empirically observe that tangent-space diffusion following Lorentz graph encoding can make the learned representations more effective for downstream multi-type PPI prediction. Specifically, when the Lorentz representations are projected into the tangent space and processed using no-return tangent diffusion, the resulting representations consistently improve prediction performance across the evaluated benchmark settings. This observation suggests that the hierarchical structural information encoded in Lorentz representations may not be fully accessible to the downstream classifier via the original decoding path. Furthermore, tangent-space diffusion can serve as a post-embedding adaptation interface for transforming such latent information into prediction-accessible signals. We refer to this phenomenon as Lorentz-to-tangent adaptation and give a mechanistic account of it in Section 4.6.

Based on this observation, we propose the Lorentz Tangent Diffusion Adapter (LTDA). In this paper, LTDA first employs a relation-specific Lorentz encoder to learn hierarchical protein representations in hyperbolic space. It then maps these Lorentz representations into the tangent space via the logarithmic map, performs lightweight graph diffusion over predefined propagation steps, and directly decodes the tangent-diffused representations for multi-type PPI prediction. Unlike manifold-closed diffusion, LTDA does not require the diffused representations to return to the Lorentz manifold prior to prediction. Instead, it treats the tangent space as a prediction-oriented adaptation interface, where latent hierarchical information encoded in Lorentz representations can be transformed into more prediction-accessible and discriminative signals. This design reflects the central hypothesis of this work: Lorentz graph encoding captures hierarchical structural information, and tangent-space diffusion at the post-embedding stage facilitates the adaptation of this information for downstream prediction.

Systematic ablation studies further demonstrate that the benefit of LTDA cannot be fully attributed to ordinary graph smoothing. Both input smoothing and output smoothing yield only limited improvements, and more complex transition priors, such as curvature-aware or geometry-aware reweighting, do not consistently outperform the uniform transition. These results indicate that the primary contribution of LTDA lies not in designing an increasingly complex diffusion kernel, but in identifying an appropriate post-embedding adaptation space for Lorentz representations. Furthermore, degree-bin analysis on SHS27K demonstrates that LTDA enhances prediction across sparse, low-degree, medium-degree, and hub-related regions, indicating that its effect is not limited to hub correction or local smoothing. Prediction-score migration and external biological evidence analysis further support the biological plausibility of LTDA-corrected protein interaction predictions.

The main contribution of this work is to introduce a post-embedding adaptation view for hyperbolic PPI prediction. Instead of only asking how to embed proteins into Lorentz space, we examine how the learned Lorentz representations should be adapted and decoded for downstream prediction. Based on this perspective, we propose LTDA, a lightweight no-return tangent diffusion adapter that projects Lorentz representations into the tangent space, performs uniform diffusion, and uses the resulting representations for multi-type PPI prediction. We further characterize, at the level of protein pairs, where LTDA improves predictions and whether the corrected pairs are biologically plausible, through degree-bin analysis, prediction-score migration, and external biological evidence analysis.

## 2. Results

### 2.1 LTDA improves multi-type PPI prediction across graph splits

We first evaluated whether post-embedding tangent-space diffusion improves multi-type PPI prediction across different graph scales and split strategies. LTDA was compared with representative PPI prediction methods, including HIGH-PPI, BaPPI, MAPE-PPI and a reproduced HI-PPI baseline, on SHS27K and SHS148K under both breadth-first search (BFS) and depth-first search (DFS) splits. These four settings jointly assess performance under different network sizes and train–test structural distributions. All reproduced results are reported as mean ± standard deviation over five independent runs using the same data splits and evaluation protocol. Differences between LTDA and the reproduced HI-PPI baseline were assessed with paired tests on the aligned test-set predictions (DeLong test for AUC; paired bootstrap for AUPR, Micro-F1 and accuracy), using one-sided tests in the a priori direction of improvement (Section 4.9); two-sided Holm–Bonferroni-adjusted p values are reported in S1 Table.

Across the four benchmark settings, LTDA achieved the best or most competitive performance among all compared methods (Table 1). The improvement was most consistent for AUPR, which is particularly relevant for multi-label PPI prediction because interaction labels are imbalanced and positive labels are sparse relative to all possible edge-type assignments. On SHS27K, LTDA improved over the reproduced HI-PPI baseline under both BFS and DFS splits. All four metrics improved significantly under both splits (one-sided p < 0.001; Table 1). Under SHS27K-BFS, LTDA achieved 77.35% Micro-F1, 83.27% AUPR, 91.77% AUC and 84.33% accuracy, compared with 74.55%, 77.16%, 89.37% and 82.08% for HI-PPI. Under SHS27K-DFS, LTDA achieved 79.49% Micro-F1, 85.76% AUPR, 93.97% AUC and 86.94% accuracy, again exceeding HI-PPI across all four metrics. These results indicate that the gain is not restricted to one graph partitioning strategy.

**Table 1.**
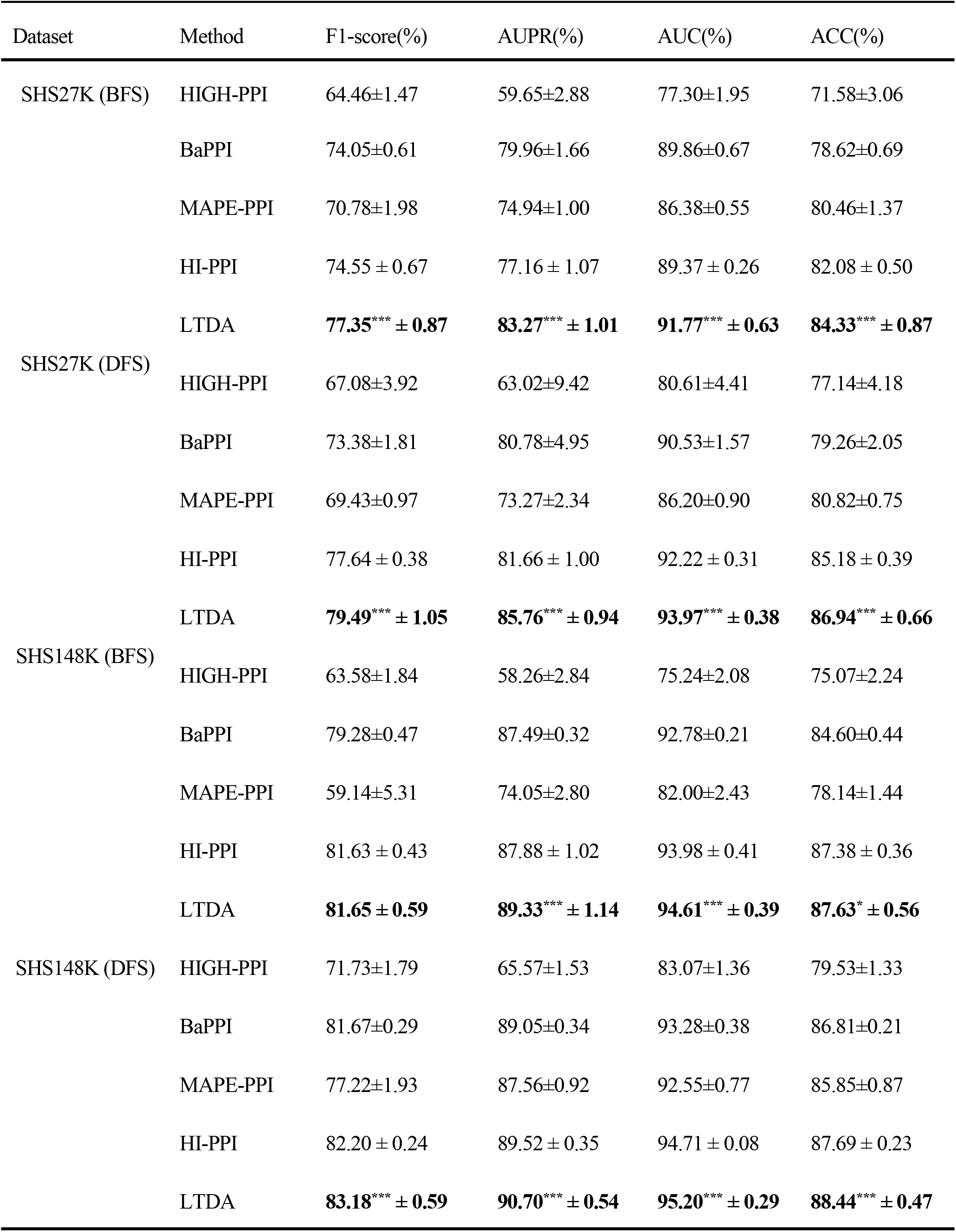
1 Prediction performance on SHS27K and SHS148K under BFS and DFS splits. Results are reported as mean ± std over five independent runs. Bold indicates the best result in each setting. Asterisks indicate significant differences from the reproduced HI-PPI baseline (*p < 0.05, **p < 0.01, ***p < 0.001; DeLong test for AUC, paired bootstrap for the remaining metrics, one-sided tests in the a priori direction of improvement; two-sided Holm–Bonferroni-adjusted p values in S1 Table).

On the larger SHS148K dataset, LTDA remained competitive and continued to improve ranking-oriented and threshold-based metrics. Under SHS148K-BFS, LTDA achieved a Micro-F1 comparable to HI-PPI, while improving AUPR, AUC and accuracy. Under SHS148K-DFS, LTDA improved HI-PPI by 0.98 percentage points in Micro-F1, 1.18 percentage points in AUPR, 0.48 percentage points in AUC and 0.75 percentage points in accuracy. Overall, compared with the reproduced HI-PPI baseline, LTDA achieved average gains of approximately 1.41 percentage points in Micro-F1, 3.21 percentage points in AUPR, 1.31 percentage points in AUC and 1.25 percentage points in accuracy across the four settings. The consistently larger improvement in AUPR suggests that LTDA enhances the model’s ability to rank positive interaction labels under class imbalance, rather than only shifting threshold-dependent decisions. All differences on SHS148K were significant except Micro-F1 under BFS, where the two methods were effectively tied (81.65% vs. 81.63%, one-sided p = 0.076).

The magnitude of improvement varied with dataset scale and split setting. The largest gains were observed on SHS27K, whereas the gains on SHS148K were more moderate but still stable, especially under DFS. This pattern is consistent with the hypothesis that tangent-space adaptation is particularly useful when the available network is smaller or structurally less redundant. In denser networks, the baseline hyperbolic encoder may already capture more useful neighbourhood information, leaving less room for post-embedding adaptation. Nevertheless, the absence of performance collapse across SHS27K and SHS148K indicates that LTDA is not over-specialized to a single dataset scale. Together, these results show that no-return tangent-space diffusion provides a robust improvement over a strong hyperbolic PPI baseline across graph splits, dataset sizes and evaluation metrics.

### 2.2 Tangent-space adaptation of Lorentz representations

We next investigated why LTDA improves prediction. The central hypothesis of this work is that Lorentz graph encoding captures hierarchical structural information, but this information may not be fully accessible to the downstream decoder unless it is adapted after embedding. LTDA addresses this post-embedding problem by projecting Lorentz representations into tangent space, applying lightweight diffusion and directly decoding the tangent-diffused representations. To test whether the observed improvement reflects a Lorentz-specific post-embedding effect rather than a generic benefit of adding diffusion, we compared the effect of the same no-return tangent diffusion operation on both the Lorentz branch and a Hyperboloid control branch.

The two branches showed markedly different responses (Fig 1). On the Lorentz branch, adding LTDA produced consistent positive gains across all four metrics under both SHS27K-DFS and SHS27K-BFS. Under DFS, the gains reached +4.69, +6.29, +3.00 and +3.98 percentage points in Micro-F1, AUPR, AUC and accuracy, respectively. Under BFS, the corresponding gains were +6.27, +7.54, +3.96 and +5.22 percentage points. In contrast, applying the same no-return tangent diffusion operation to the Hyperboloid branch did not produce a comparable pattern. The Hyperboloid branch decreased under DFS and showed only near-zero changes under BFS. Thus, the improvement cannot be explained simply by assuming that diffusion benefits any hyperbolic representation. Instead, the result suggests a selective interaction between Lorentz representations and no-return tangent-space adaptation.

**Fig 1.**
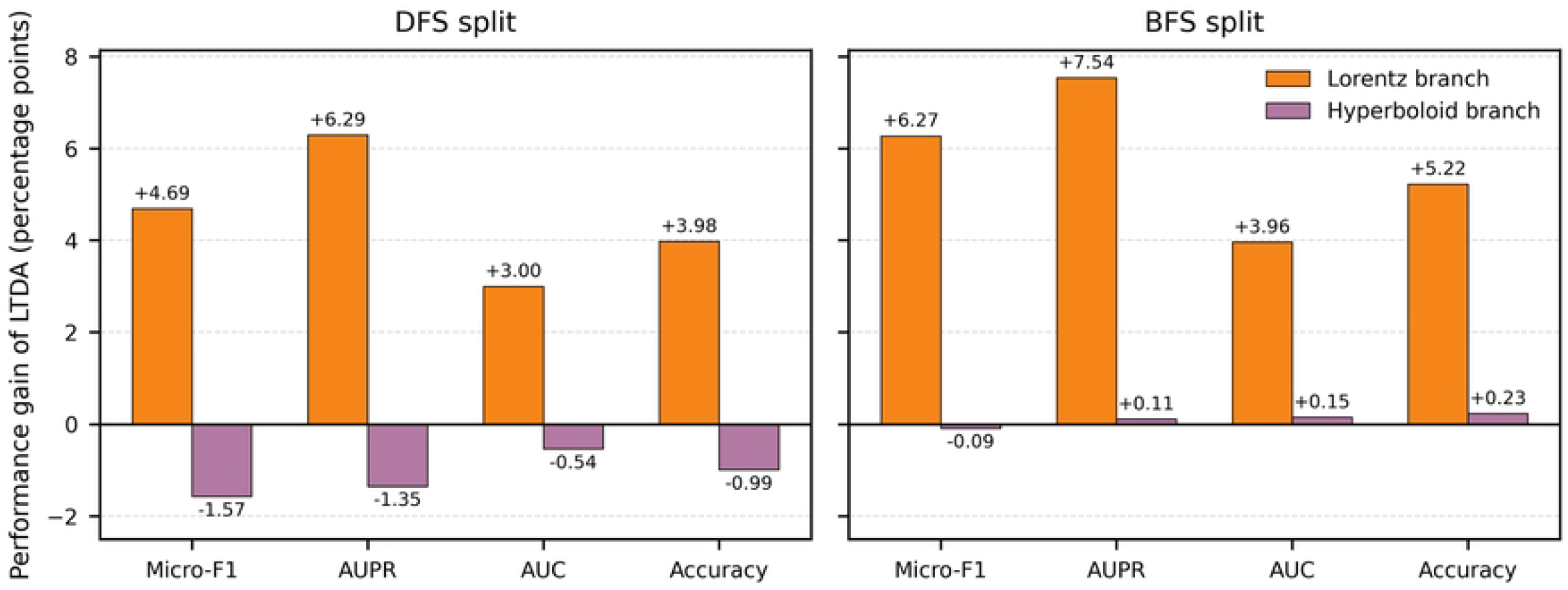
Effectiveness of LTDA and its dependence on the hyperbolic branch. The bar plot shows the performance gain of LTDA over the corresponding backbone without LTDA on the SHS27K DFS and BFS splits, measured in percentage points. The gain is defined as *Δ* = P*erf*o*rmance*_*+LTDA*_*− Performance*_*w/*o*LTDA*_. LTDA brings consistent positive gains to the Lorentz branch.

We then asked whether LTDA is equivalent to ordinary graph smoothing. Two controls were considered: input smoothing, which applies graph diffusion to Euclidean input features before model encoding, and output smoothing, which applies diffusion to prediction outputs after decoding. These controls represent two common ways in which graph diffusion may improve performance without invoking a post-embedding hyperbolic mechanism. To provide a strong comparison, we used the metric-wise best result among input and output smoothing as the ordinary-smoothing reference. Although ordinary smoothing improved performance to some extent, its gains were consistently smaller than those of LTDA (Fig 2). Under DFS, LTDA provided additional gains over the best ordinary smoothing reference across all four metrics. The same trend was observed under BFS, where LTDA again outperformed ordinary smoothing by a clear margin. These results indicate that the benefit of LTDA is not adequately explained by smoothing raw input features or post-processing prediction outputs.

**Fig 2.**
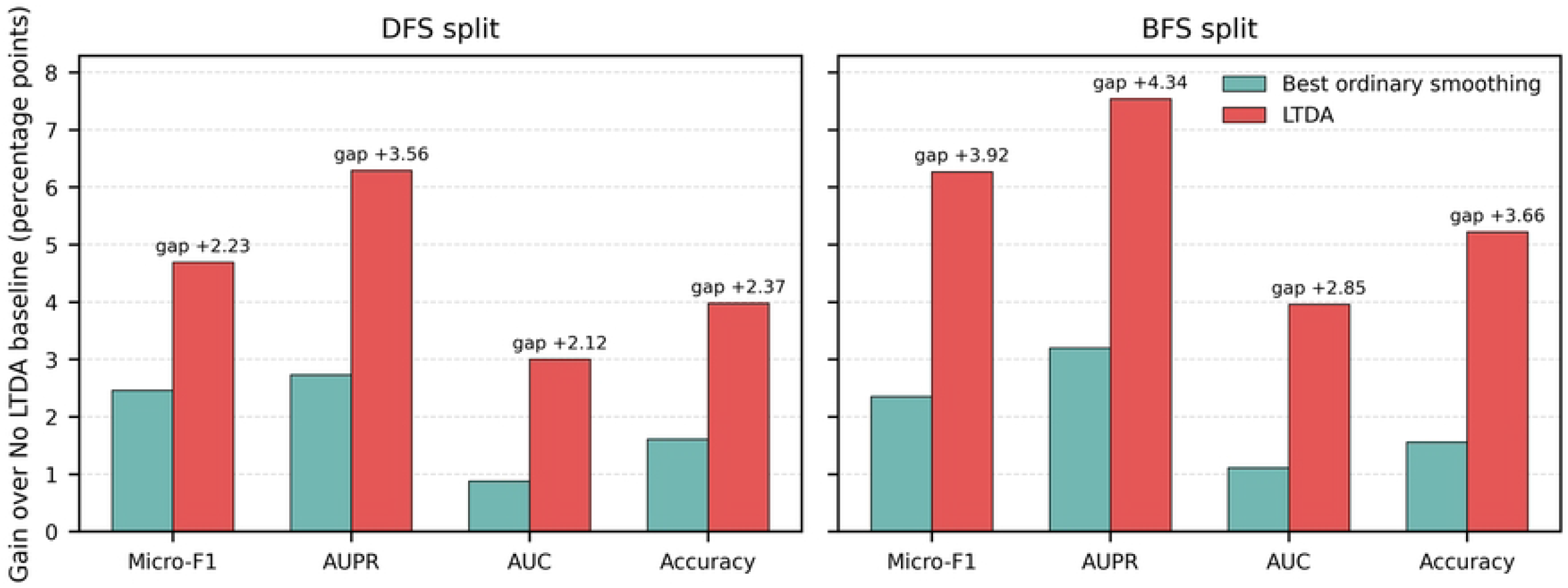
Comparison between LTDA and ordinary smoothing. The bar plot shows the performance gains of the metric-wise best ordinary smoothing variant and LTDA over the No-LTDA baseline, measured in percentage points. Best ordinary smoothing denotes the better result selected from input smoothing and output smoothing for each evaluation metric, and is used as a stronger metric-wise reference among the tested ordinary smoothing controls. The gap indicates the additional improvement of LTDA over this reference. Although ordinary smoothing brings certain gains, LTDA achieves further improvements under both DFS and BFS splits, suggesting that the observed benefit of LTDA is not adequately explained by input feature smoothing or prediction post-processing alone, but is more consistent with a Lorentz-to-tangent adaptation effect (Section 4.6).

A further question is whether tangent-diffused representations should be mapped back to the manifold before prediction. We therefore compared no-return decoding with return-manifold decoding after tangent diffusion (Fig 3). As analysed in Section 4.6, the two variants are equivalent up to the numerical realization of the exponential–logarithmic round trip and the parameterization of the readout, so this comparison quantifies the cost of a redundant round trip rather than a difference in information. Significance was assessed with the DeLong test for AUC and a paired bootstrap over test pairs for the remaining metrics, applied to each run (Section 4.9). For the Lorentz branch, no-return decoding was better in every metric and under both splits. Under SHS27K-BFS the improvement was significant for all four metrics: AUC 94.24 ± 0.30 vs. 93.79 ± 0.18 (p < 0.001), Micro-F1 83.39 ± 0.32 vs. 82.27 ± 0.10, i.e. 1.12 percentage points (p = 0.004), accuracy 87.15 ± 0.18 vs. 86.34 ± 0.21, i.e. 0.81 points (p = 0.002), and AUPR 88.87 ± 0.68 vs. 88.12 ± 0.32, i.e. 0.75 points (p = 0.026). Under SHS27K-DFS the AUC (95.04 ± 0.17 vs. 94.73 ± 0.06, p = 0.004), AUPR (89.18 ± 0.19 vs. 88.60 ± 0.15, p = 0.029) and accuracy (87.90 ± 0.36 vs. 87.28 ± 0.20, p = 0.018) differences were significant, whereas the Micro-F1 difference (80.85 ± 0.44 vs. 80.23 ± 0.18, 0.63 points, p = 0.085) was in the same direction but not significant. The Hyperboloid control branch showed the opposite response: return-manifold decoding was significantly better in AUC under both splits (p = 0.017 and p < 0.001), in AUPR under BFS (0.59 points, p = 0.004) and, under DFS, in accuracy (0.96 points, p = 0.002), Micro-F1 (1.12 points, p = 0.013) and AUPR (0.73 points, p = 0.024), whereas the two remaining comparisons of that branch were not significant. Because the two branches differ in the placement of nonlinearity and in how curvature enters the diffusion kernel (Section 4.6), this contrast is best described as implementation-dependent; within the Lorentz branch evaluated here, it supports the minimal, no-return form of tangent-space decoding.

**Fig 3.**
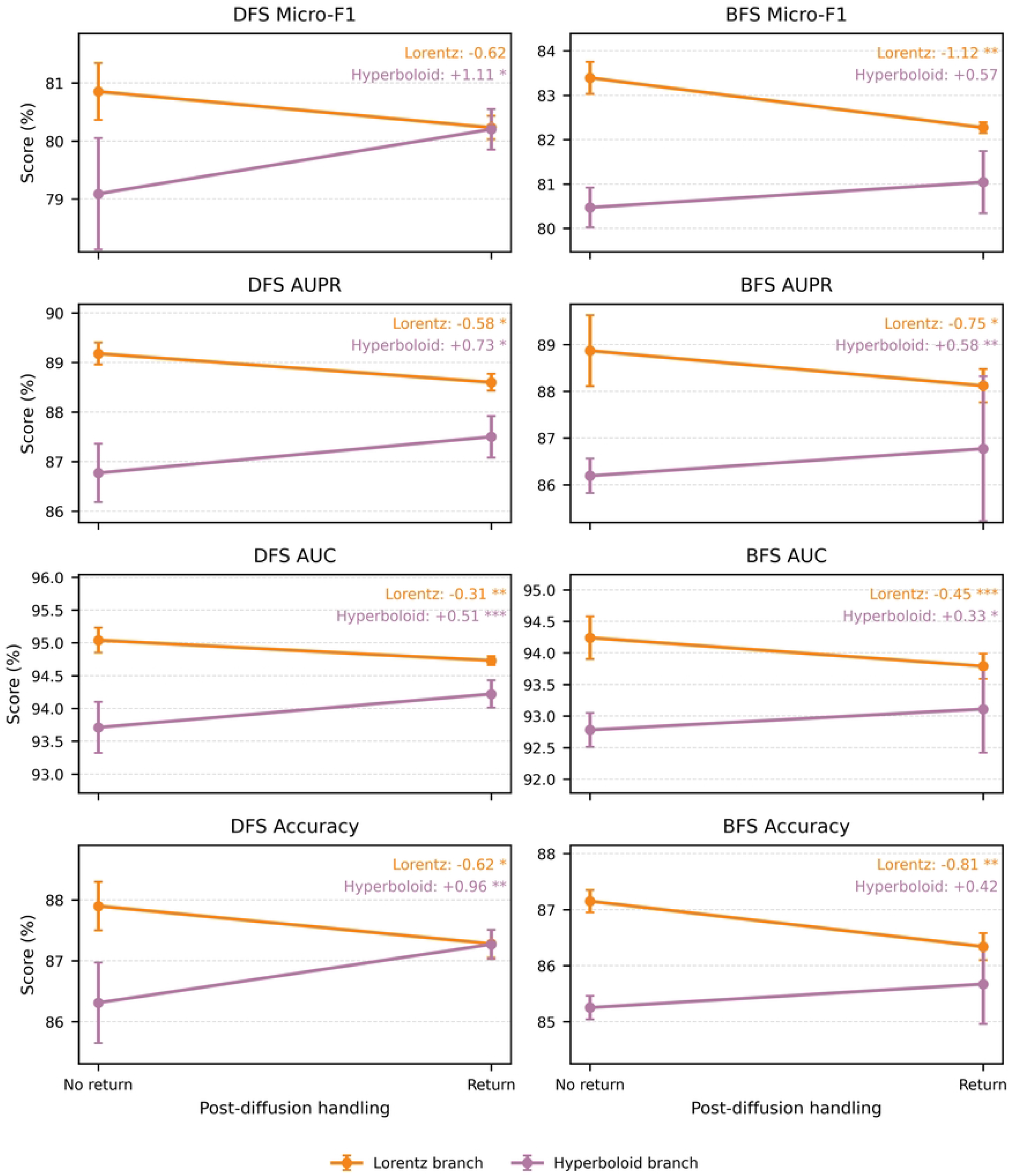
Comparison between return-manifold and no-return decoding after tangent diffusion.

Micro-F1, AUPR, AUC and ACC of the Lorentz and Hyperboloid branches under the SHS27K DFS and BFS splits. No-return decoding predicts directly from the tangent-diffused representations, whereas return-manifold decoding maps them back to the hyperbolic manifold first; in exact arithmetic the two are equivalent up to the numerical realization of the exponential–logarithmic round trip and the readout parameterization (Section 4.6). Under the evaluated settings the Lorentz branch favours no-return decoding, whereas the Hyperboloid control branch shows the opposite trend, reflecting differences between the two hyperbolic implementations rather than a universal property of hyperbolic representations. Error bars denote the standard deviation over five runs; annotated values are Return − No return (percentage points), and asterisks mark significant differences between the two strategies (*p < 0.05, **p < 0.01, ***p < 0.001; Section 4.9; two-sided Holm–Bonferroni-adjusted p values in S1 Table).

Finally, we examined whether more complex diffusion transitions are necessary. Uniform diffusion was compared with curvature-aware and geometry-aware transition priors under SHS27K-BFS and SHS27K-DFS. These priors did not provide stable advantages over the uniform transition. Although small metric-specific improvements were observed in a few cases, they were not consistent across splits or evaluation metrics, and in several settings the complex priors reduced AUPR or AUC. Since AUPR is especially informative in class-imbalanced multi-label PPI prediction, a small improvement in Micro-F1 at the cost of ranking performance does not constitute a stable advantage. Therefore, the gain of LTDA appears to arise primarily from the post-embedding Lorentz-to-tangent adaptation interface, rather than from increasingly complex transition-prior reweighting. Uniform diffusion provides a lightweight and stable default for adapting Lorentz representations into prediction-accessible signals.

Together, these ablation results support a Lorentz-to-tangent adaptation effect (Section 4.6). LTDA is not merely adding diffusion to a graph model, smoothing input features, smoothing prediction outputs or designing a more complex transition kernel. Instead, its key role is to place a simple diffusion operation at a specific point in the modelling pipeline: after Lorentz graph encoding and before downstream decoding. This post-embedding location allows hierarchical information encoded in Lorentz representations to become more accessible for multi-type PPI prediction.

### 2.3 LTDA improves predictions across sparse and hub-related regions

PPI networks contain proteins with highly uneven connectivity. Some proteins have no observed neighbours in the training graph, whereas hub proteins participate in many interactions and often belong to multiple functional modules. A performance gain averaged over all test edges may therefore hide distinct behaviours across sparse, low-degree and hub-related regions. To determine where LTDA improves prediction, we performed degree-bin analysis on SHS27K under both BFS and DFS splits. For each test protein pair, the pair-level degree was defined as the maximum training-graph degree of its two endpoint proteins. Test pairs were grouped into five bins: degree 0, 1–2, 3–9, 10–29 and ≥30.

LTDA improved the Lorentz branch across all degree bins and all four metrics under both split settings (Fig 4). Under DFS, positive gains were observed not only in medium- and high-degree regions, but also in sparse and low-degree regions. AUPR increased in every bin, with particularly large gains for hub-related pairs. Under BFS, the improvement was again broadly distributed across degree bins. Notably, the degree-0 bin, in which both endpoint proteins have no observed neighbours in the training graph, also showed substantial improvement. This finding is important because degree-0 test pairs cannot benefit from ordinary neighbourhood memorization in the training graph. The improvement in this bin suggests that LTDA is not simply exploiting high-degree proteins or correcting hub-related bias.

**Fig 4.**
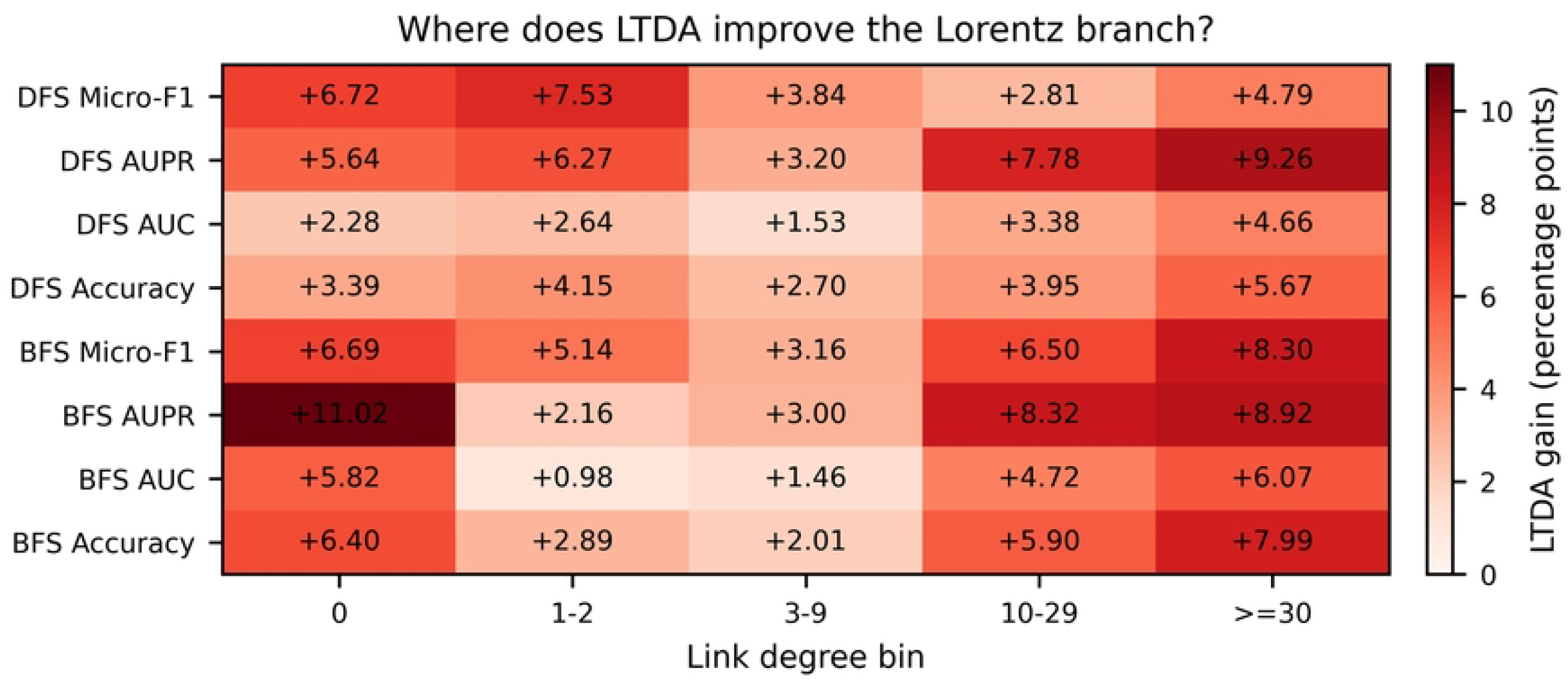
Performance improvements of LTDA over the Lorentz branch across different degree bins.

Test edges are grouped according to the maximum degree of their two endpoint proteins in the training graph, i.e., *d*(*u,v*) = max(*d*_*u*,_*d*_*v*_). The horizontal axis denotes the link degree bins, including 0, 1–2, 3–9, 10–29 and ≥30. The heatmap shows the performance gains of LTDA over the Lorentz baseline on the SHS27K DFS and BFS splits in terms of Micro-F1, AUPR, AUC, and ACC, measured in percentage points. Positive gains are observed across all degree bins and all evaluation metrics, suggesting that the benefit associated with LTDA is not limited to hub-related interactions, but can also be observed in extremely sparse and low-degree regions of the training graph. These observations provide additional support for the Lorentz-to-tangent adaptation effect under different structural conditions. The numbers of test edges in each degree bin are reported in Table 2.

**Table 2.** The numbers of test edges in each degree bin.

| edges | bfs | dfs |
| --- | --- | --- |
| [0, 1) | 299 | 199 |
| [1, 3) | 148 | 174 |
| [3, 10) | 363 | 417 |
| [10, 30) | 399 | 362 |
| [30, $1 \times 10^8$) | 319 | 378 |

The degree-bin pattern provides two insights. First, LTDA improves prediction in structurally complex regions. Hub-related interactions are not necessarily easier to predict, because hub proteins may participate in diverse biological processes and have multiple potential interaction types. The gains observed in the ≥30 bin suggest that tangent-space adaptation helps the model use Lorentz representations in regions with complex local topology. Second, LTDA also improves low-connectivity and sparse regions, including degree-0 and 1–2 bins. This argues against the explanation that LTDA functions only as a hub-correction or local-smoothing mechanism. Instead, the gains are distributed across different structural regimes of the PPI network.

These results complement the ablation analyses. If LTDA were merely smoothing local graph neighbourhoods, its effects would be expected to concentrate in regions with enough observed neighbours for propagation to be informative. Instead, the observed improvements extend from sparse regions to hub-related regions. Therefore, degree-bin analysis supports the interpretation that LTDA improves the usability of Lorentz representations more generally, rather than only correcting one specific topological subgroup of protein pairs. The numbers of test edges in each degree bin are presented in Table 2, so as to facilitate the reproducibility of the analytical results.

### 2.4 LTDA reshapes prediction scores around the decision boundary

We further examined how LTDA changes model predictions at the score level. This analysis was performed on SHS27K-BFS using predictions averaged over five independent runs. The Lorentz model without LTDA was treated as the pre-adaptation baseline, and the Lorentz model with uniform tangent diffusion was treated as the LTDA model. A fixed decision threshold of 0.5 was used to identify label-level decision changes.

LTDA increased the separation between positive and negative labels (Fig 5a). Before applying LTDA, the mean prediction score of positive labels was 0.717 and that of negative labels was 0.197, corresponding to a positive-negative margin of 0.520. After applying LTDA, the mean positive-label score increased to 0.830, whereas the mean negative-label score decreased to 0.141. The margin therefore increased to 0.689. This pattern indicates that LTDA does not simply shift all prediction scores upward. Instead, it increases scores for positive labels while suppressing scores for negative labels, producing a more discriminative score distribution.

**Fig 5.**
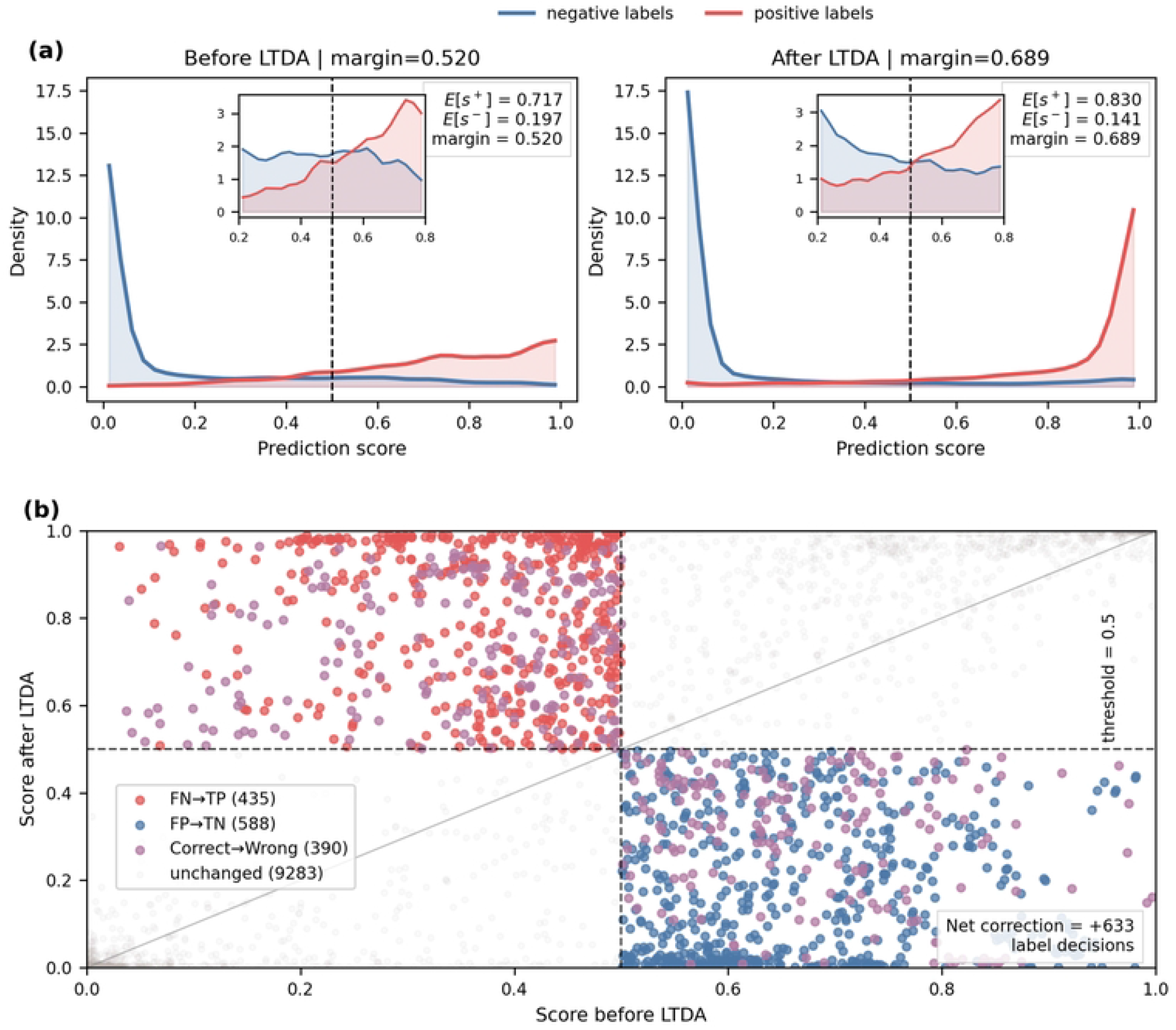
Prediction-score analysis of tangent-space adaptation. (a) Prediction-score distributions of positive and negative labels before and after LTDA on the SHS27K-BFS split. “Before LTDA” denotes the Lorentz baseline without LTDA, whereas “After LTDA” denotes the Lorentz model equipped with uniform tangent diffusion. Prediction scores are averaged over five independent runs. The prediction margin is defined as *E*[*s*^+^]−*E*[*s*^−^]. LTDA increases the mean positive-label score from 0.717 to 0.830 and decreases the mean negative-label score from 0.197 to 0.141, enlarging the margin from 0.520 to 0.689. The inset zooms into the score range [0.2, 0.8] around the decision boundary. (b) Label-level score migration before and after LTDA. Each point denotes one edge-label prediction. The dashed lines indicate the fixed decision threshold 0.5, and the diagonal line denotes unchanged prediction score. LTDA converts 435 false negatives into true positives and 588 false positives into true negatives, while converting 390 previously correct label predictions into incorrect ones, resulting in a net correction of +633 label-level decisions. Correct→Wrong denotes label-level predictions that are correct before LTDA but incorrect afterwards.

The label-level migration pattern further supports this interpretation (Fig 5b). Many false-negative labels before LTDA moved above the decision threshold after LTDA and became true positives, while many false-positive labels moved below the threshold and became true negatives. Specifically, LTDA converted 435 false-negative labels into true positives and 588 false-positive labels into true negatives. At the same time, it introduced 390 cases that were correct before LTDA but became incorrect after LTDA, resulting in a net correction of +633 label-level decisions. Therefore, the improvement is not driven by indiscriminate score inflation. Rather, LTDA tends to increase under-confident positive predictions and suppress over-confident false-positive predictions.

This score-level analysis provides a mechanistic view of the performance gains observed in the benchmark experiments. The improved AUPR in the main results is consistent with better separation between positive and negative labels, while the increased number of corrected label-level decisions explains the improvements in threshold-dependent metrics such as Micro-F1 and accuracy. However, the score-migration analysis should be interpreted as empirical evidence for the prediction behaviour of LTDA, not as a proof of the adaptation mechanism formalized in Section 4.6. It shows that tangent-space adaptation changes the decision landscape in a direction that is consistent with improved multi-label PPI classification.

### 2.5 External evidence supports LTDA-corrected interactions

The preceding analyses show that LTDA improves benchmark metrics and reshapes prediction scores. We next asked whether the corrected predictions are biologically plausible. Because the benchmark labels are derived from existing PPI annotations and external biological resources are incomplete, this analysis was designed as supporting evidence rather than direct experimental validation. We focused on label-level false-negative decisions corrected by LTDA, denoted as FN→TP cases, on the SHS27K-BFS split. Prediction scores were averaged over five independent runs, and the same fixed decision threshold used in the main evaluation was applied to identify decision changes. A high-confidence corrected subset was further defined by requiring an LTDA score of at least 0.8 and a score gain of at least 0.3 over the Lorentz baseline.

To provide a controlled background, we sampled degree-matched unlabeled protein pairs from non-benchmark pairs. Matching was performed using the same pair-level degree definition as in the degree-bin analysis, based on the maximum training-graph degree of the two endpoint proteins. Separate matched backgrounds were generated for all FN→TP cases and for the high-confidence corrected subset. Since unobserved protein pairs in incomplete PPI networks cannot be assumed to be true negatives, these background pairs were treated only as unlabeled controls.

Four categories of external biological evidence were considered: curated physical interactions from BioGRID, co-complex membership from CORUM, shared Reactome events or pathways, and functional overlap based on GOA human annotations. To reduce overly broad functional matches, IEA annotations were excluded and GO terms annotated to more than 5% of proteins were removed. Strict GO support was defined as sharing at least one non-broad GO Biological Process or Molecular Function term.

LTDA-corrected FN→TP cases were strongly enriched for external evidence compared with degree-matched unlabeled controls (Table 3). Among all FN→TP cases, 45.52% were supported by at least one evidence source under the strict GO definition, whereas only 4.83% of the degree-matched background pairs had such support. The high-confidence corrected subset showed a similar pattern, with 47.76% of corrected pairs supported by at least one evidence source compared with 3.67% of its matched background. Reactome provided particularly strong pathway-level support: 31.26% of all FN→TP cases and 36.33% of high-confidence corrected cases shared at least one Reactome event or pathway, whereas the corresponding backgrounds showed 0.46% and 0.00% Reactome support, respectively. Evidence from BioGRID physical interactions, CORUM complexes and strict GO functional overlap also showed higher hit rates in LTDA-corrected cases than in matched controls.

**Table 3.**
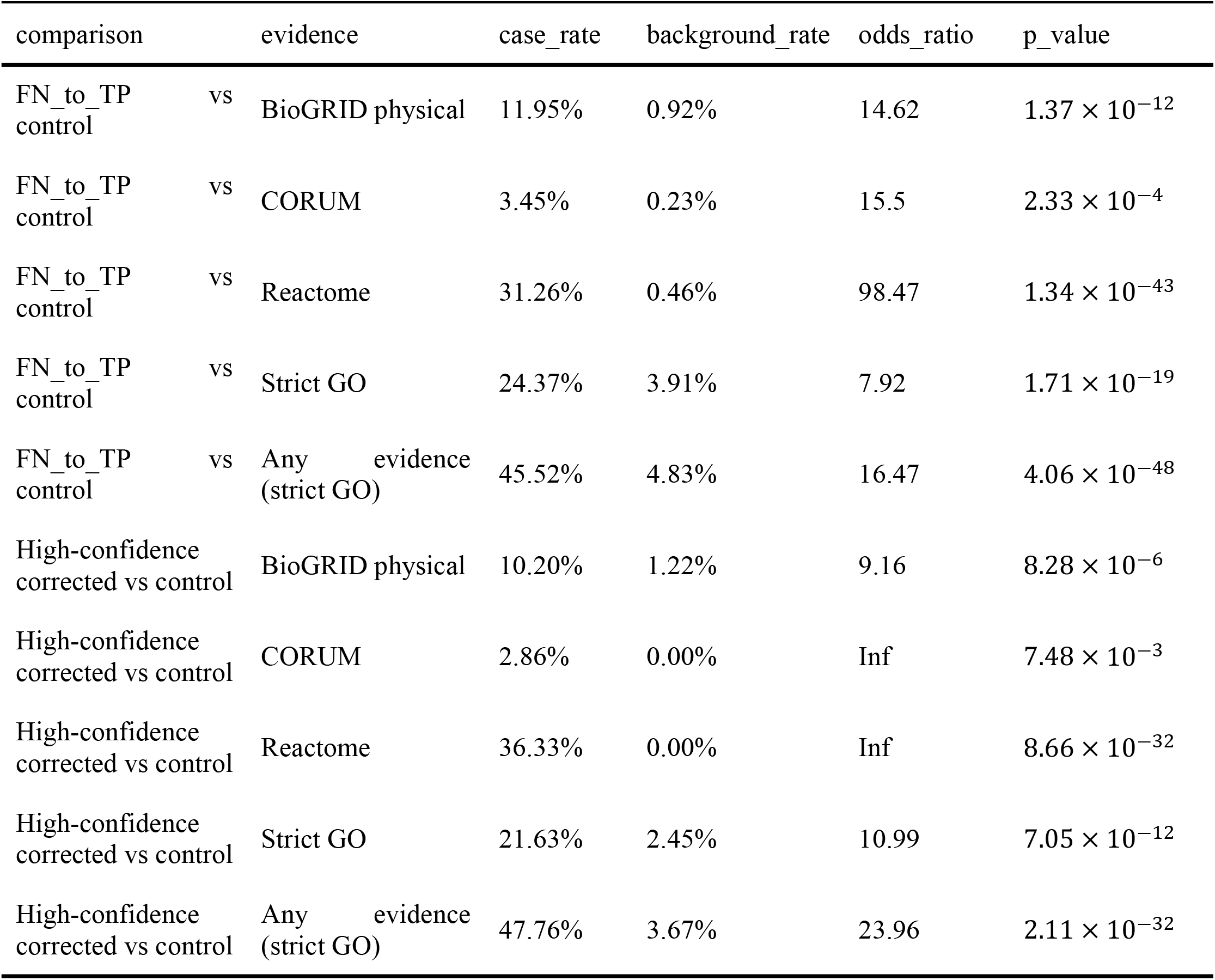
External biological evidence enrichment of LTDA-corrected interactions. Evidence hit rates are compared between LTDA-corrected cases and degree-matched unlabeled background controls on the SHS27K-BFS split. Here, control denotes degree-matched unlabeled protein-pair background pairs sampled from non-benchmark protein pairs, with the same training-graph degree-bin distribution as the corresponding LTDA-corrected group. For the FN→TP group, the control was matched to all FN→TP cases; for the high-confidence corrected group, a separate control was matched to the high-confidence subset. Any evidence (strict GO) denotes support from BioGRID physical, CORUM, Reactome, or strict GO functional overlap. Odds ratios and p-values were computed using one-sided Fisher’s exact test. Infinite odds ratios occur when no matched control pair has the corresponding evidence; Fisher’s exact test was still computed using the original contingency table. External evidence was evaluated at the protein-pair level and should be interpreted as biological plausibility rather than direct validation of each STRING action-specific label.

These enrichments indicate that a substantial fraction of LTDA-recovered positive decisions correspond to protein pairs with independent biological support. The evidence is strongest at the protein-pair or pathway level, which is expected because most external resources do not provide STRING action-specific labels for every interaction type. Therefore, absence of external support should not be interpreted as biological implausibility, and presence of external support should not be interpreted as direct validation of each predicted interaction label. Instead, the analysis supports the biological plausibility of the corrected protein pairs and provides an independent sanity check that LTDA does not merely improve benchmark metrics by exploiting statistical artefacts.

Overall, the Results establish a coherent evidence chain. LTDA improves multi-type PPI prediction across two datasets and two graph split strategies. Ablation analyses show that the improvement is better explained by post-embedding Lorentz-to-tangent adaptation than by ordinary smoothing, return-manifold decoding or complex transition priors. Degree-bin analysis shows that the improvement is distributed across sparse, low-degree and hub-related regions. Score-migration analysis shows that LTDA increases positive-negative score separation and corrects more label-level decisions than it disrupts. Finally, external biological evidence analysis indicates that LTDA-corrected predictions are enriched for independent biological support. Together, these results suggest that the post-embedding stage is a critical and previously underexplored component of hyperbolic PPI representation learning.

## 3. Discussion

Accurate prediction of protein–protein interactions remains challenging because PPI networks are incomplete, noisy, class-imbalanced, and structurally heterogeneous. In this study, we proposed LTDA, a Lorentz Tangent Diffusion Adapter, to improve hyperbolic multi-type PPI prediction by introducing tangent-space diffusion at the post-embedding stage of Lorentz representation learning. Unlike conventional graph diffusion methods that operate on Euclidean input features or final prediction outputs, LTDA acts after Lorentz graph encoding, where hierarchical structural information has already been embedded but may not be fully accessible to the downstream classifier. Results on SHS27K and SHS148K show that LTDA improves prediction performance across different benchmark settings, and the ablation studies suggest that these gains are more consistent with an empirically observed Lorentz-to-tangent adaptation effect than with generic graph smoothing or complex transition-prior reweighting.

A central observation of this work is that no-return tangent diffusion provides consistent gains for the Lorentz implementation branch under the evaluated settings, and Section 4.6 gives a mechanistic account of why this form is the appropriate one. In the SHS27K BFS and DFS ablations, applying tangent-space diffusion after Lorentz graph encoding improves multiple evaluation metrics, whereas applying the same operation to the Hyperboloid control branch does not yield stable gains. This contrast suggests that the benefit of LTDA is unlikely to be explained simply by assuming that diffusion improves any hyperbolic representation. Instead, under our implementation and benchmark settings, Lorentz representations appear to contain prediction-relevant structural information that can become more accessible after projection and propagation in the tangent space. The return-manifold comparison shows that decoding the diffused representation directly is at least as good as, and in most metrics better than, applying an additional exponential–logarithmic round trip before the readout; as shown in Section 4.6, the two paths are equivalent up to the numerical realization of that round trip and the parameterization of the readout, so the no-return form is the minimal realization of tangent-space decoding. Therefore, these findings should be interpreted as empirical evidence for a geometry-dependent post-embedding decoding pattern, rather than as a general theoretical claim about all hyperbolic representations.

The smoothing-control and transition-prior ablations further clarify the nature of LTDA. Ordinary input smoothing and output smoothing provide only limited improvements, suggesting that LTDA is not adequately explained as an input preprocessing or output post-processing technique. Curvature-aware and geometric-aware transitions also do not show stable advantages over uniform diffusion. These observations indicate that increasing the complexity of the diffusion kernel is not the main factor associated with the observed performance improvement. Instead, the results support the interpretation that a simple tangent-space diffusion interface can serve as a stable post-embedding adaptation space for Lorentz representations. This is particularly meaningful for PPI networks, where hub proteins, missing edges, and experimental noise may make complex transition priors sensitive to local structural bias.

The structural, prediction-level, and biological analyses provide complementary support for this empirical interpretation. Degree-bin analysis shows that LTDA improves the Lorentz branch across sparse, medium-degree, and hub-related regions, suggesting that its effect is not limited to hub correction or local smoothing. Score-migration analysis further shows that LTDA corrects many false-negative and false-positive label-level decisions, with corrected decisions exceeding newly introduced errors. Together with the enlarged positive-negative prediction margin, this suggests that LTDA changes the decision distribution in a more discriminative direction rather than simply inflating all prediction scores. In addition, LTDA-corrected FN →TP cases show stronger external biological support than degree-matched unlabeled controls, using evidence from BioGRID physical interactions, CORUM complexes, Reactome pathways, and stringent GO functional overlap. These results support the biological plausibility of LTDA-corrected predictions at the protein-pair level, although they should not be interpreted as direct experimental validation of each STRING action-specific label.

Several limitations should be acknowledged. First, SHS27K and SHS148K are derived from STRING, and the benchmarks may still reflect biases in existing PPI annotations, even though independent resources are used for biological evidence analysis. Second, the analysis of biological evidence primarily offers support at the protein-pair level or pathway level and does not directly validate each predicted interaction type. Third, this work focuses on computational validation, and wet-lab experiments are required to confirm high-confidence candidate interactions that are not recorded in databases. Fourth, the contrast between the Lorentz and Hyperboloid branches reflects differences between two hyperbolic GNN paradigms — the placement of nonlinearities, the treatment of curvature and the parameterization of the readout — rather than a difference in the tangent-space construction itself. The mechanistic analysis in Section 4.6 explains why the effect should be expected to be model-specific; quantifying it across curvature values and hyperbolic architectures is left for future work.

In future work, LTDA can be extended by integrating stronger protein language models or structure foundation models to provide richer initial protein representations [38–40]. Label-aware or pathway-aware tangent diffusion may further improve the discrimination of interaction types such as catalysis, activation, inhibition, and binding. LTDA may also be applied to other hierarchical and scale-free biological networks, including gene regulatory networks, drug–target interaction networks, and disease–gene association networks. Overall, this study suggests that the post-embedding stage is a critical but underexplored component of hyperbolic PPI representation learning. By projecting Lorentz representations into the tangent space, performing lightweight uniform diffusion, and directly using the tangent-diffused representation for prediction, LTDA provides an effective and lightweight strategy for predicting PPI types.

## 4. Methods

### 4.1 Datasets

We conducted experiments on two public human protein–protein interaction benchmark datasets, namely SHS27K and SHS148K. Both datasets are derived from the Homo sapiens subset of the STRING database and have been widely utilized in previous studies for evaluating multi-type PPI prediction models [25]. SHS27K contains 1,690 proteins and 12,517 protein–protein interaction records, whereas SHS148K contains 5,189 proteins and 44,488 protein–protein interaction records. The two datasets exhibit substantial differences in network scale, protein coverage, and the number of interactions, thereby enabling the evaluation of model performance and generalization across varying graph sizes and interaction densities.

Throughout this paper, multi-type refers to the number of interaction categories (seven STRING action types in this study), whereas multi-label refers to the formal setting in which each protein pair may carry several types simultaneously; in this sense multi-type prediction is implemented as a multi-label classification problem, and we use interaction type and label interchangeably. In terms of task formulation, we model PPI prediction as a multi-label classification task. For each candidate protein pair, the model is required to determine whether it belongs to one or more interaction types. Based on the STRING action labels provided in the benchmark, we consider seven common PPI types, including reaction, binding, post-translational modification, activation, inhibition, catalysis, and expression. Since the same protein pair may participate in multiple biological interaction processes, the multi-label prediction setting better reflects the coexistence of different interaction relationships in real PPI networks. In addition, the distributions of different interaction types are imbalanced, with some minority interaction types containing relatively limited samples. Therefore, we report Micro-F1, AUPR, AUC, and ACC to comprehensively evaluate model performance under both overall prediction and class-imbalanced conditions [26, 27].

To evaluate model stability and generalization under different graph partitioning conditions, we adopt two split strategies, Breadth-First Search (BFS) and Depth-First Search (DFS), to construct the training and test sets. For each dataset, approximately 20% of PPI edges are selected as the held-out test set according to the corresponding BFS or DFS strategy, while the remaining interaction edges are used for model training. The test set is used only for final performance evaluation and is not involved in model training. BFS and DFS splits result in distinct train–test structural distributions, enabling the assessment of model performance under different network sampling conditions. Compared with a single random split, employing both BFS and DFS settings aids in evaluating the model’s adaptability to different protein coverage patterns, local connectivity structures, and potential distribution shifts. This partitioning protocol is consistent with prior PPI prediction studies such as HI-PPI.

For input features, we use the publicly available benchmark-provided preprocessed protein sequence and structure-related models, and follow the same feature construction pipeline as previous studies without introducing additional features. The initial representation of each protein comprises sequence descriptors and structural encodings. Sequence features include pseudo amino acid composition (PAAC), composition/transition/distribution (CTD), and global positional information, whereas structural features are extracted from protein structure-related representations using a pretrained structural encoding module [28–30]. Finally, the sequence and structural features are concatenated to form the initial node representations in the PPI graph.

Overall, we evaluate the proposed model under four standard experimental settings: SHS27K-BFS, SHS27K-DFS, SHS148K-BFS, and SHS148K-DFS. This setup enables the evaluation of model performance on PPI networks of varying scales, as well as the evaluation of model stability, robustness, and generalization under different graph partitioning strategies.

### 4.2. Problem Formulation

Let *G* = (*V,E,X,Y*) denote a multi-label protein--protein interaction graph, where 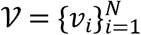 is the set of proteins, ℰis the set of protein pairs, 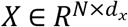 is the initial protein feature matrix, and *Y* ∈ {0,1}^|*E*|×*C*^ is the multi-label interaction matrix. Here, *N* denotes the number of proteins, *d*_x_ denotes the dimension of the initial protein feature, and *C* denotes the number of interaction types. In this work, *C* = 7, corresponding to reaction, binding, post-translational modification, activation, inhibition, catalysis, and expression.

For a candidate protein pair (*v*_*i*,_*v*_*j*_), its ground-truth multi-label vector is defined as

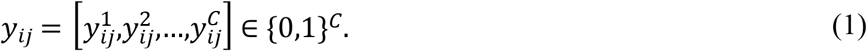

Here, *y*_*ij*_ denotes the ground-truth interaction label vector of protein pair (*v*_*i*_,*v*_*j*_), and 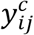 denotes the binary label for the *c*-th interaction type. Specifically, 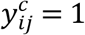 indicates that protein pair (*v*_*i*_,*v*_*j*_) has the *c*-th type of interaction, whereas 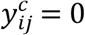 indicates that this interaction type is absent. Since one protein pair may have multiple interaction types simultaneously, *y*_*ij*_ is a multi-label vector rather than a single-class label.

The model outputs a logit vector

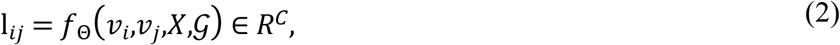

where *f*_Θ_(·) denotes the proposed PPI prediction model parameterized by Θ, and 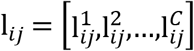 denotes the predicted logit vector for protein pair *v*_*i*_,*v*_*j*_. Each 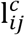 is the unnormalized prediction score for the *c*-th interaction type.

The predicted multi-label probability vector is

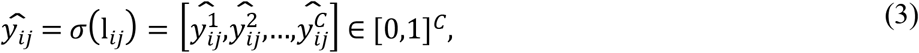

where σ(·) denotes the element-wise sigmoid function, and 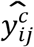 denotes the predicted probability that protein pair (*v*_*i*,_*v*_*j*_) belongs to the *c*-th interaction type.

### 4.3 Overall Architecture

As shown in Fig 6, the proposed framework consists of four main components: Lorentz feature mapping, relation-specific Lorentz graph encoding, Lorentz Tangent Diffusion Adapter, and gated pair-wise interaction prediction.

**Fig 6.**
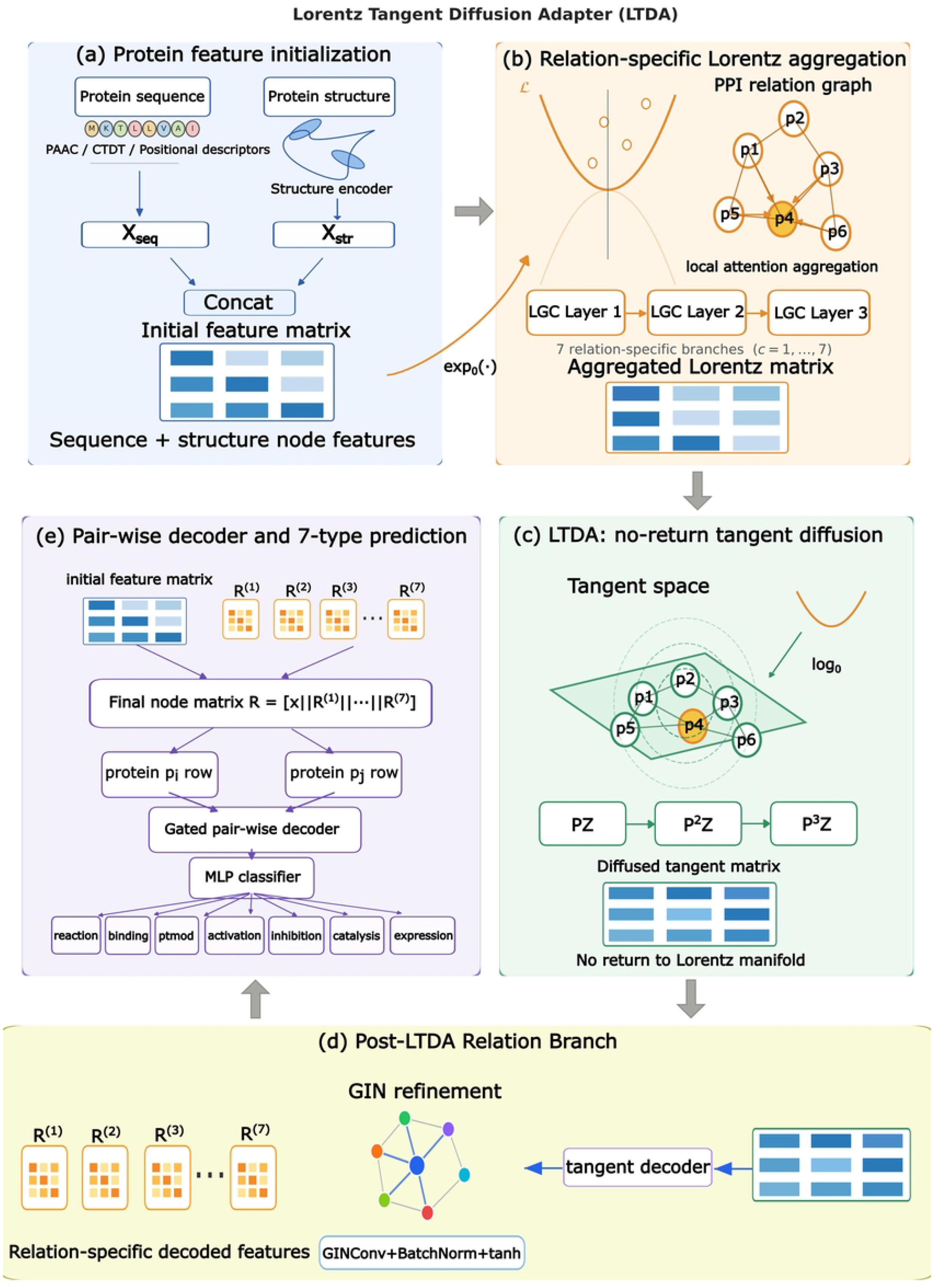
Overall architecture of the proposed Lorentz Tangent Diffusion Adapter (LTDA) for multi-type protein– protein interaction prediction. Protein sequence descriptors and structure-derived encodings are concatenated to form the initial protein feature matrix. The protein representations are then mapped into the Lorentz manifold and processed by relation-specific Lorentz graph aggregation. LTDA projects the learned Lorentz representations into the tangent space, performs uniform multi-scale diffusion, and directly decodes the tangent-diffused representations without returning them to the manifold. The post-LTDA relation branches are further refined by relation-specific GIN modules. Finally, the original features and all relation-specific decoded features are concatenated and fed into a gated pair-wise decoder followed by an MLP classifier to predict seven PPI types.

Given the initial protein feature matrix *X*, each protein feature is first mapped into the Lorentz manifold. Then, for each interaction type *c* ∈ {1,…,*C*}, an independent relation-specific Lorentz branch with *L* = 3 graph convolution layers are used to learn relation-aware protein representations. Following Lorentz encoding, LTDA maps the Lorentz representations to the tangent space, performs uniform tangent diffusion, and directly decodes the tangent-diffused representations without remapping them back to the Lorentz manifold. Finally, all relation-specific decoded representations are concatenated with the original feature representation and fed into a gated pair-wise decoder.

For protein *v*_*i*_, the final node representation is

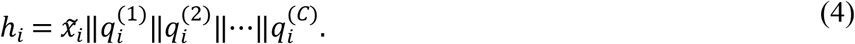

Here, *h*_*i*_ denotes the final representation of protein *v*_*i*_, 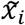 denotes the original protein feature after zero-time-coordinate augmentation, 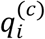 denotes the decoded representation of protein *v*_*i*_ from the *c*-th relation branch, and ‖ denotes vector concatenation. This formulation integrates the original protein feature and all relation-specific representations into a unified node representation for downstream pair-wise prediction.

### 4.4 Relation-specific Lorentz Graph Encoding

To model the hierarchical and hub-heavy structure of PPI networks, we employ the Lorentz model of hyperbolic space. Let 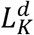 denote a *d*-dimensional Lorentz manifold embedded in *R*^*d*+1^:

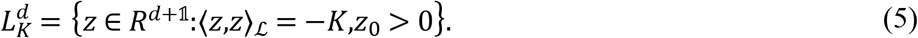

Here, *z* = [*z*_0,_*z*_1_,…,*z*_*d*_] denotes a point on the Lorentz manifold, *z*_0_ is the Lorentz time coordinate, and *z*_1_, …,*z*_*d*_ are the spatial coordinates. *K* > 0 is a curvature-related radius parameter, and *z*_0_ > 0 ensures that the representation lies on the upper sheet of the Lorentz manifold.

The Lorentzian inner product is defined as

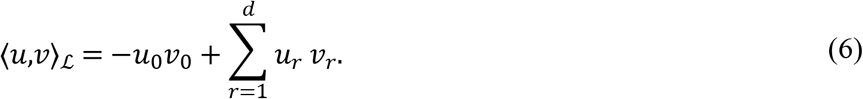

Here, *u,v* ∈ *R*^*d*+1^ are two Lorentz-space vectors, *u*_0_ and *v*_0_ are their time coordinates, and *u*_*r*_, *v*_*r*_ for *r* = 1,…,*d* are their spatial coordinates. This inner product defines the geometry of the Lorentz manifold.

For each protein feature x_*i*_, we first add a zero time coordinate:

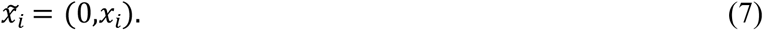

Here, 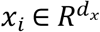 is the initial feature vector of protein *v*_*i*_, and 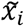 denotes the augmented feature after adding a zero time coordinate. This operation makes the input compatible with the Lorentz representation format.

The initial Lorentz representation is obtained by projecting 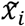 to the tangent space at the origin and applying the exponential map:

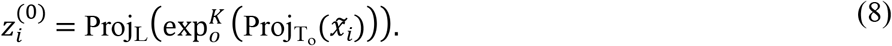

Here,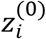 denotes the initial Lorentz representation of protein *v*_*i*_, o denotes the Lorentz origin, 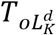 denotes the tangent space at the origin, 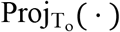 projects the augmented feature to the tangent space, 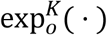 maps the tangent vector to the Lorentz manifold, and Proj_L_(·) enforces the Lorentz manifold constraint for numerical stability.

For each interaction type *c*, an independent Lorentz branch is used. Let 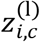 denote the Lorentz representation of protein *v*_*i*_ in branch *c* at layer *l*. The initial representation of each relation branch is

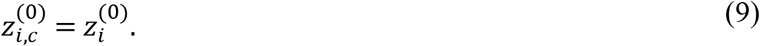

Here, 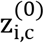 is the initial representation of protein *v*_*i*_ in the *c*-th relation branch, and it is initialized from the shared Lorentz representation 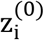.

At the l*-*th Lorentz graph convolution layer, a Lorentz transformation is first applied:

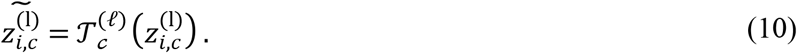

Here, 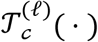 denotes the Lorentz transformation in the *c*-th relation branch at layer l, and 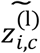 denotes the transformed representation of protein *v*_*i*_.

Since local attention aggregation is enabled in the main configuration, the neighbor aggregation is written as

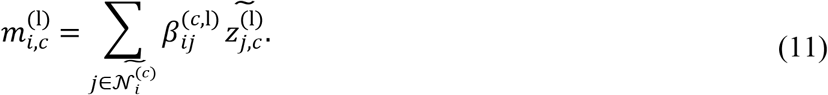

Here, 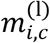 denotes the aggregated representation of protein *v*_*i*_ at layer l in relation branch *c*, 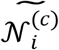 denotes the self-loop-augmented neighbor set of protein *v*_*i*_ in relation graph *c*, and 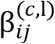 denotes the local attention coefficient measuring how much information protein *v*_*j*_ contributes to protein *v*_*i*_. The summation aggregates information from the neighbors of *v*_*i*_ under the *c*-th interaction type.

The aggregated representation is then normalized to satisfy the Lorentz constraint:

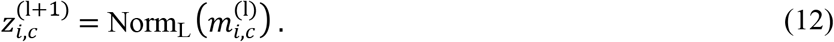

Here, Norm_L_ (·) denotes Lorentz normalization, which projects or normalizes the aggregated representation so that it remains compatible with the Lorentz manifold constraint. 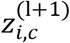 is the updated Lorentz representation of protein *v*_*i*_ for the next layer.

After *L* = 3 Lorentz graph convolution layers, the final Lorentz representation matrix of relation branch *c* is

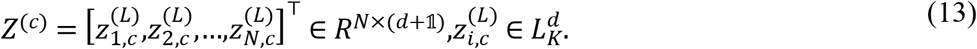

Here, *Z*^*(c*)^ denotes the final Lorentz representation matrix of all proteins in the *c*-th relation branch. Its *i*-th row is 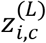, which is the final Lorentz representation of protein *v*_*i*_ after *L* = 3 Lorentz graph convolution layers.

### 4.5 Lorentz Tangent Diffusion Adapter

Although Lorentz graph encoding can represent hierarchical PPI structures, the learned hyperbolic embeddings may still contain latent structural information that is not fully accessible to the downstream prediction decoder. To make these latent signals accessible to the downstream prediction decoder, we propose the Lorentz Tangent Diffusion Adapter.

LTDA first maps Lorentz representations into the tangent space, then performs graph diffusion in the tangent space, and finally directly employs the tangent-diffused representation for prediction. In the primary LTDA configuration, uniform tangent diffusion and no-return decoding are adopted. Specifically, the diffused representation is not mapped back to the Lorentz manifold prior to prediction.

Given the Lorentz representation matrix *Z*^*(c*)^ from the *c*-th relation branch, LTDA first maps it to the tangent space at the Lorentz origin:

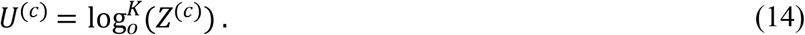

Here, *U*^*(c*)^ denotes the tangent-space representation obtained from the Lorentz representation matrix *Z*^*(c*)^, and 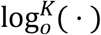 denotes the logarithmic map at the Lorentz origin o. This operation transforms Lorentz representations into a locally Euclidean tangent space.

Since the first coordinate corresponds to the Lorentz time component, LTDA uses only the spatial tangent component for subsequent diffusion. Let *π*_*s*p_(·) denote the operation that removes the time coordinate and keeps the spatial coordinates. The tangent-space representation matrix is

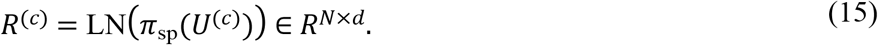

Here, *R*^(*c*)^ denotes the normalized spatial tangent representation matrix for relation branch *c, π*_sp_(*U*^*(c*)^) extracts the spatial coordinates from *U*^*(c*)^, and LN(·) denotes layer normalization. Each row 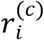 of *R*^(*c*)^ is the tangent-space representation of protein *v*_*i*_.

In the main configuration, LTDA adopts a uniform tangent transition. Let 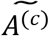 denote the adjacency matrix of relation *c* after symmetrization and self-loop addition. The row-normalized transition matrix is

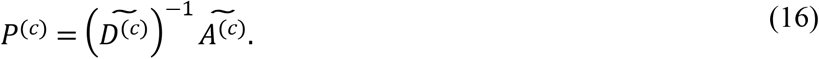

Here, P^*(c*)^ denotes the uniform transition matrix used for tangent diffusion in relation branch *c*, 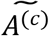 denotes the self-loop-augmented adjacency matrix, and *D*^*(c*)^ denotes the corresponding degree matrix. Each element 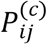 controls how much information protein *v*_*j*_ contributes to protein *v*_*i*_ during diffusion.

The degree matrix is defined as

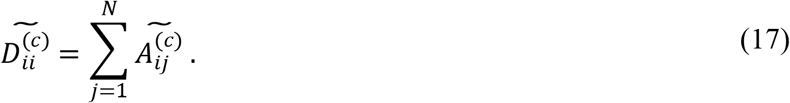

Here, 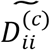 is the degree of protein *v*_*i*_ in the self-loop-augmented relation graph *c*, computed as the sum of the *i*-th row of 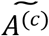. This normalization makes the transition weights in each row sum to one.

For diffusion scale *s*, the propagated tangent representation is

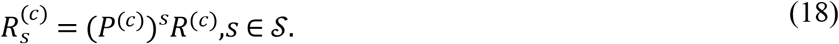

Here, 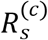 denotes the tangent representation after *s-*step graph diffusion, (*P*^*(c*)^) ^*s*^ means applying the transition matrix *s* times, and *S* is the set of diffusion scales. When *s* = 1, the representation mainly aggregates one-hop neighborhood information; when *s* > 1, it incorporates higher-order neighborhood information.

In the main experiments, the diffusion scale set is fixed as

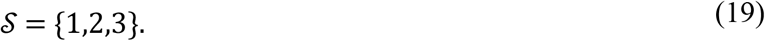

Here, *S* contains the diffusion depths used by LTDA. The setting {1,2,3} means that LTDA fuses one-hop, two-hop, and three-hop tangent diffusion representations.

The multi-scale diffused representation is obtained by learnable scale fusion:

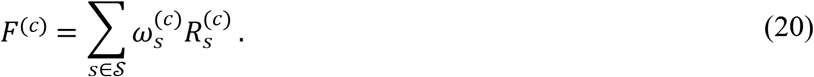

Here, *F*^*(c*)^ denotes the fused multi-scale tangent-diffused representation for relation branch *c*, and 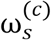 is the learnable fusion weight for diffusion scale *s*. This equation combines different propagation scales into one representation.

The scale weight is defined as

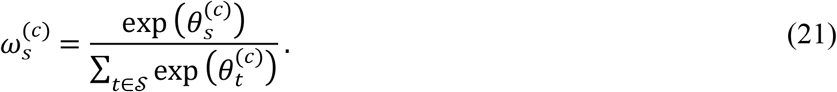

Here, 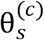 is the learnable scale-fusion logit for scale *s* in relation branch *c*, and the softmax normalization ensures that the weights over all scales sum to one.

To preserve the original tangent representation and reduce excessive smoothing, LTDA further applies a degree-aware residual gate:

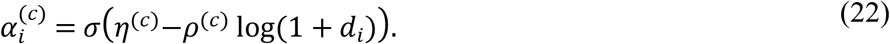

Here, 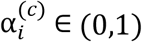 is the residual diffusion strength for protein *v*_*i*_ in relation branch *c, d*_*i*_ is the degree of protein *v*_*i*_, *η*^*(c*)^ is a learnable residual-strength parameter, and *ρ*^*(c*)^ ≥ 0 is a learnable degree attenuation coefficient. The term *l*o*g(*1 + *d*_*i*_) makes the gate degree-aware while avoiding numerical instability for low-degree or zero-degree nodes.

The final residual-gated tangent representation of protein *v*_*i*_ in relation branch *c* is

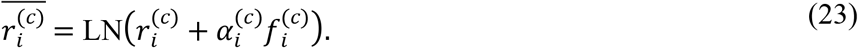

Here, 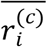 denotes the final tangent representation after residual-gated diffusion, 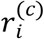 is the original tangent representation of protein *v*_*i*_, and 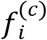 is the *i*-th row of the multi-scale diffused representation *F*^*(c*)^. The term 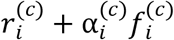 preserves the original representation while injecting diffused neighborhood information.

The corresponding matrix form is denoted as

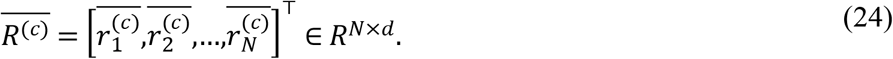

Here, 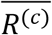 denotes the final residual-gated tangent representation matrix for all proteins in relation branch *c. F*^*(c*)^ denotes the multi-scale diffused representation, whereas 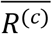 denotes the residual-gated tangent representation finally used for no-return decoding.

Because the main configuration sets *ltda*_return_manifold = false, LTDA directly decodes the tangent-diffused representation without mapping it back to the Lorentz manifold:

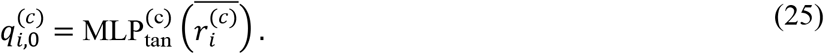

Here, 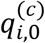 denotes the initial decoded tangent representation of protein *v*_*i*_ from relation branch *c*, and ML 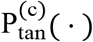 is the tangent-space decoder for the *c*-th relation branch. No exponential map is applied after tangent diffusion.

The decoded representations are further processed by a relation-specific Euclidean GIN layer:

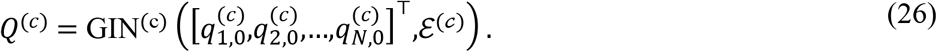

Here, GIN^(c)^ (·) denotes the relation-specific Euclidean GIN module, and *E*^*(c)*^ denotes the edge set of the *c*-th relation graph used by this GIN layer. This step further refines the no-return tangent-decoded representations in Euclidean space.

The output of the relation-specific GIN layer is

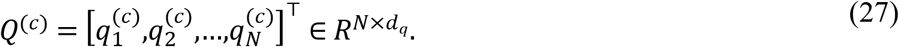

Here, *Q*^*(c*)^ denotes the final decoded representation matrix of relation branch *c*, and 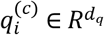 is the final relation-specific representation of protein *v*_*i*_. This no-return design conceptualizes the tangent space as a prediction-oriented adaptation interface. Lorentz encoding captures latent hierarchical structure, while tangent diffusion and no-return decoding convert these structure into linearly readable predictive signals. The following subsection explains why the tangent space is the appropriate space for this adaptation and why no-return decoding is the minimal form of it.

### 4.6 A mechanistic account of tangent-space decoding

Graph diffusion is a linear operator on a vector space. For a row-stochastic transition matrix **P** and a node-feature matrix **H**, the filtered signal ∑_*k*_ *w*_*k*_**P**^*k*^**H** is defined only when **H** belongs to a vector space, because the filter requires addition and scalar multiplication of features. The Lorentz model is the hyperboloid *L* = {**x** ∈ ℝ^d*+1*^ :⟨**x**,**x**⟩_*L*_ = −*K*, x_0_ > 0}, which is a nonlinear submanifold: the coordinate-wise average of two points on *L* generally does not satisfy the constraint, so neighbourhood averaging cannot be applied to Lorentz representations directly. The tangent space at the origin, *T*_**o**_*L*, is the canonical linearization of *L*: the logarithmic map log_**o**_ is a smooth bijection from *L* onto *T*_**o**_*L*≅*ℝ*^*d*^, and its inverse *e*x*p*_**o**_ restores the manifold constraint exactly. Placing diffusion in *T*_**o**_*L* is therefore not a modelling shortcut but the exact setting in which the diffusion operator is defined: a manifold-closed propagation is the same computation followed by a nonlinear map that only re-imposes the constraint. Because the readout in LTDA is linear, the pipeline “filter then linear readout” is also exactly the graph-signal-processing construction ∑_*k*_ *w*_*k*_**P**^*k*^**H**, which is not available on the manifold itself.

In Lorentz coordinates the time coordinate is not a free parameter: it is fixed by the spatial coordinates through the manifold constraint, 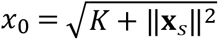 (with the constants of the respective parameterization). The manifold therefore carries *d* free degrees of freedom, and the additional channel is a nonlinear function of the radius. Two consequences follow for the post-embedding stage. First, every tangent vector at the origin has an identically zero time component, so the tangent representation of a point is fully determined by its spatial part and no information is discarded when the time channel is removed before diffusion (Eq. 15). Second, in Lorentz coordinates the time coordinate grows like cosh of the radial distance whereas the spatial coordinates grow like sinh, so the time channel is dominated by a radial, almost common-mode term that is shared by all nodes at a similar depth. Normalizing and filtering the spatial tangent components therefore operates on the discriminative part of the representation and keeps the linear readout in a space where features are comparable across nodes; re-introducing a zero time channel after diffusion, as the returning variant does, adds a channel that is constant and hence carries no information.

The returning variant applies *e*x*p*_**o**_ to the diffused tangent representation and then log_**o**_ again before the linear readout. On the tangent space at the origin, *e*x*p*_**o**_ ° log_**o**_ is the identity map, so in exact arithmetic the two variants provide the classifier with the same tangent representation; they differ in the additional nonlinear maps placed between diffusion and classification, in the numerical safeguards these maps contain (the exponential map clips the tangent norm, a saturating operation that compresses representations with large radius, i.e. nodes lying deep in the hierarchy or with high degree), and in the parameterization of the readout that follows. Returning to the manifold therefore adds no information, and whatever perturbation it introduces acts directly on the representation that the diffusion has just produced. The no-return form removes this redundant composite and hands the filtered representation to the classifier in the same linear space in which it was computed. This also explains why the differences we observe between the two variants are small, metric-dependent and always in favour of the no-return form rather than qualitative (Section 2.2), and why we treat no-return decoding as the canonical realization of tangent-space adaptation rather than as a separate modelling device.

The Lorentz and Hyperboloid branches used in our ablation correspond to two different hyperbolic GNN paradigms rather than to two parameterizations of the same model. The Lorentz branch follows the HyboNet formulation: linear maps and nonlinearities act in the tangent space at the origin and the manifold constraint is restored after each layer. The Hyperboloid branch follows the Chami formulation: nonlinearities act in the tangent space of the current point, and curvature is a learnable parameter that enters the exponential and logarithmic maps themselves; in our implementation the diffusion kernel is coupled to this learnable curvature on the Hyperboloid branch, whereas the corresponding maps of the Lorentz branch are evaluated at unit curvature at the origin. The LTDA block therefore receives representations that differ in scale, in the placement of nonlinearity and in how curvature enters the geometry of the maps, and because the kernel normalizes and filters its input, a difference between branches is expected rather than anomalous. This is why we present the effect as a property of a specific hyperbolic encoder together with a specific adaptation path, not as a property of hyperbolic representations in general, and why characterizing the benefit as a function of curvature and of the placement of nonlinearities is a natural direction for future work.

### 4.7 Gated Pair-wise Interaction Decoder

After relation-specific encoding and LTDA-based tangent decoding, the final node representation of protein *v*_*i*_ is constructed as

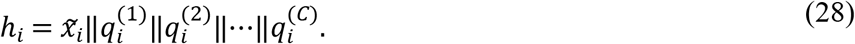

Here, *h*_*i*_ is the final representation of protein *v*_*i*_, 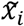 is the augmented original feature representation, and 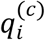 is the decoded representation from relation branch *c*. The concatenation integrates original protein information with all relation-specific LTDA representations.

For a candidate protein pair (*v*_*i*,_*v*_*j*_), the basic pair-wise interaction feature is computed by element-wise multiplication:

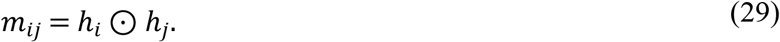

Here, *m*_*ij*_ denotes the multiplicative interaction feature of protein pair (*v*_*i*,_*v*_*j*_), and ⊙ denotes element-wise multiplication. This operation captures dimension-wise compatibility between the two protein representations.

A gated interaction network is then employed to adaptively select informative pair-wise signals:

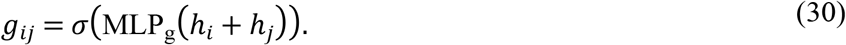

Here, *g*_*ij*_ denotes the learned interaction gate for protein pair (*v*_*i*,_*v*_*j*_), MLP_g_ (·) denotes the gate network, and *h*_*i*_ + *h*_*j*_ provides an order-invariant pair summary. The sigmoid function constrains each gate value to the range [0,1].

The preliminary gated pair representation is

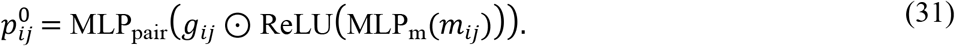

Here, MLP_m_ (·) extracts hidden interaction features from *m*_*ij*_, ReLU (·) introduces nonlinearity, *g*_*ij*_ adaptively filters the hidden interaction features, and MLP_pair_ (·) produces the preliminary pair representation 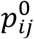.

When degree information is available, a degree-aware pair gate is further applied. The degree-related pair feature is defined as

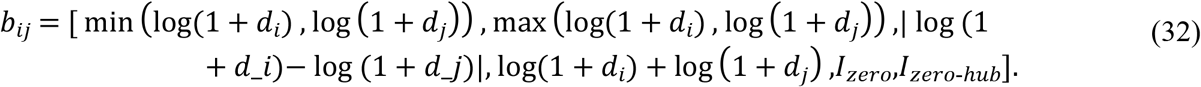

Here, *b*_*ij*_ denotes the degree-related feature vector for protein pair (*v*_*i*,_*v*_*j*_), *d*_*i*_ and *d*_*j*_ are the degrees of proteins *v*_*i*_ and *v*_*j*_, respectively. The first four terms describe the minimum, maximum, difference, and sum of the log-degrees of the two endpoints. *I*_*zer*o_ indicates whether at least one endpoint has zero degree, and *I*_*zer*o-*hub*_ indicates whether one endpoint has zero degree while the other endpoint is a high-degree protein.

The degree-aware modulation vector is

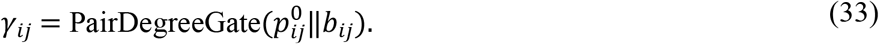

Here, *γ*_*ij*_ denotes the degree-aware modulation vector, PairDegreeGate (·) is a learnable gate network, and 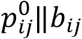 concatenates the preliminary pair representation with the degree-related pair feature.

The final pair representation is

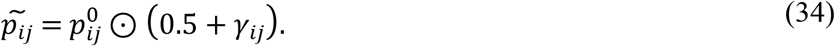

Here, 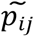 denotes the final degree-modulated pair representation. The factor 0.*5* + *γ*_*ij*_ adaptively rescales the preliminary pair representation according to degree-related structural information. When degree information is unavailable, we set 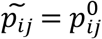.

Finally, the predicted logits are obtained by an MLP classifier:

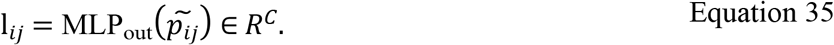

Here, *l*_*ij*_ denotes the predicted logit vector for protein pair (*v*_*i*,_*v*_*j*_), and MLP_out_(·) is the final classifier.

The *c*-th element 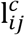 is the logit score for the *c*-th interaction type.

The predicted multi-label probability is

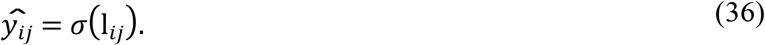

Here, 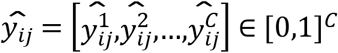 denotes the predicted probability vector for protein pair (*v*_*i*_,*v*_*j*_), and 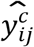 is the predicted probability that this protein pair has the *c*-th interaction type. Since this is a multi-label task, each interaction type is predicted independently through the sigmoid function.

In implementation, zero-degree endpoint proteins can be further stabilized by anchor-based representation imputation. This component is kept fixed across the default LTDA model and all ablation variants.

### 4.8 Loss Function and Prediction

The model is trained as a multi-label classification model. For each protein pair (*v*_*i*,_*v*_*j*_), the ground-truth label vector is *y*_*ij*_ ∈ {0,1}^*C*^, and the model outputs logits *l*_*ij*_ ∈ *R*^*C*^. In implementation, we use BCEWithLogitsLoss, i.e., binary cross-entropy applied directly to the model logits, which internally combines the sigmoid function with binary cross-entropy in a numerically stable manner:

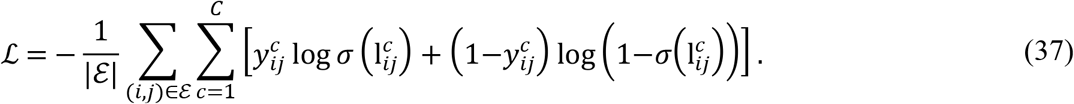

Here, *L* denotes the binary cross-entropy loss, *E*is the set of protein pairs used for training, *C* is the number of interaction types, 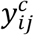 is the ground-truth label, and 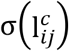 is the predicted probability for the *c*-th interaction type. This loss treats each interaction type as an independent binary classification task.

When positive class weighting is used, the weighted loss is

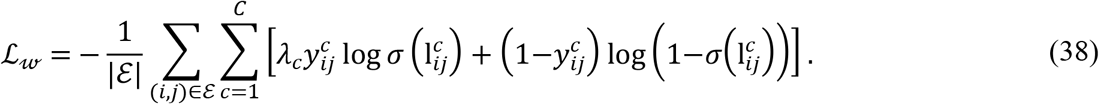

Here, *L*_w_ denotes the class-weighted binary cross-entropy loss, and *λ*_*c*_ is the positive class weight for the *c*-th interaction type. A larger *λ*_*c*_ gives higher penalty to positive samples of class *c*, which can help handle class imbalance.

During inference, prediction probabilities are obtained as

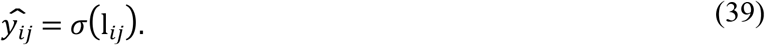

Here, *l*_*ij*_ is the output logit vector of the model, and 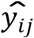 is the predicted probability vector after the sigmoid function.

Unless otherwise specified, a fixed threshold of 0.5 is used for binary prediction:

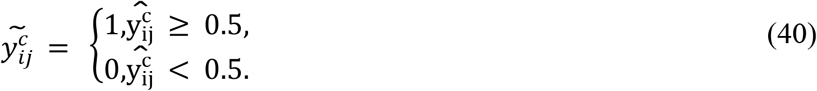

Here, 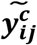 denotes the final binary prediction for the *c*-th interaction type. If the predicted probability 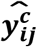 is not smaller than 0.5, the interaction type is predicted as present; otherwise, it is predicted as absent. The model is evaluated using Micro-F1, AUPR, AUC, and ACC.

### 4.9 Statistical analysis

We assessed the statistical significance of the difference between LTDA and the strongest competing method on the aligned per-pair predictions of the test sets. The direction of the expected improvement is fixed by the study design, so one-sided p values are reported for the contrasts stated a priori (LTDA over the reproduced HI-PPI baseline; no-return over return-manifold decoding); for the Hyperboloid control, where the observed differences run in the opposite direction, two-sided p values are reported instead. For each of the five runs we applied the DeLong test for two correlated ROC curves to AUC, and a paired bootstrap over test protein pairs (10,000 resamples, identical resampling indices for both methods) to AUPR, Micro-F1 and accuracy; we report the median p value across the five runs. Run-level differences were additionally summarized with a paired t-test across runs. Two-sided Holm–Bonferroni-adjusted p values, applied within each metric across the four settings of a family, are provided in S1 Table.

## Author contributions

Q.Y. and X.C. conceived and designed the study. Q.Y. and Z.X. performed the experiments and data analysis. Q.Y. developed the LTDA framework and performed the computational modeling. Q.Y., X.C. and X.X. wrote the manuscript with input from all authors. All authors discussed the results and approved the final version.

## Competing interests

The authors declare no competing interests.

## Acknowledgements

This work was supported by the National Natural Science Foundation of China [Grant Nos. 32260154 and 62562041].

## Data and code availability

The source code for the LTDA framework is available at https://github.com/one-chun/LTDA. All other data supporting the findings of this study are available within the article and its supplementary information files, or from the corresponding author upon reasonable request.

## Correspondence

Correspondence and requests for materials should be addressed to Xiang Cheng and Xuan Xiao.

## Supporting information

S1 Table. Two-sided Holm–Bonferroni-adjusted p values for the statistical comparisons in the main text. The table lists the uncorrected per-run p values reported in the main text together with their two-sided Holm– Bonferroni-adjusted counterparts for the comparison of LTDA with the reproduced HI-PPI baseline (Table 1) and for the comparison of no-return with return-manifold decoding (Fig 3).

## Notes

### Competing Interest Statement

The authors have declared no competing interest.

